# Human Gene Expression Variability and its Dependence on Methylation and Aging

**DOI:** 10.1101/500785

**Authors:** Nasser Bashkeel, Theodore J. Perkins, Mads Kærn, Jonathan M. Lee

## Abstract

**Background:** Phenotypic variability of human populations is partly the result of gene polymorphism and differential gene expression. As such, understanding the molecular basis for diversity requires identifying genes with both high and low population expression variance and identifying the mechanisms underlying their expression control. Key issues remain unanswered with respect to expression variability in human populations. The role of gene methylation as well as the contribution that age, sex and tissue-specific factors have on expression variability are not well understood.

**Results:** Here we used a novel method that accounts for sampling error to classify human genes based on their expression variability in normal human breast and brain tissues. We find that high expression variability is almost exclusively unimodal, indicating that variance is not the result of segregation into distinct expression states. Genes with high expression variability differ markedly between tissues and we find that genes with high population expression variability are likely to have age-, but not sex-dependent expression. Lastly, we find that methylation likely has a key role in controlling expression variability insofar as genes with low expression variability are likely to be non-methylated.

**Conclusions:** We conclude that gene expression variability in the human population is likely to be important in tissue development and identity, methylation, and in natural biological aging. The expression variability of a gene is an important functional characteristic of the gene itself and the classification of a gene as one with Hyper-Variability or Hypo-Variability in a human population or in a specific tissue should be useful in the identification of important genes that functionally regulate development or disease.

## Background

Within the last decade, many studies have established that gene expression patterns vary between individuals, across tissue types[1], and within isogenic cells in a homogenous environment[2]. These differences in gene expression lead to phenotypic variability across a population. Differential gene expression gene expression is typically detected by analyzing expression data from a population of samples in two or more genetic or phenotypic states, for example a cancerous and non-cancerous sample or between two different individuals. Various differential gene expression algorithms, such as edgeR and DESeq, are then used to identify genes whose expression mean differs significantly between the states. While differential co-expression analyses have successfully been used to identify novel disease-related genes[3], the statistical methods used in these analyses consider gene expression variance within the sample population as a component of the statistical significance estimate. However, expression variability within populations has been emerging as an informative metric of cell state an informative metric of a phenotypic state, particularly as it relates to human disease[4, 5].

There are several sources of expression variability in a population. The first are polymorphisms that contribute, both genetically and epigenetically, to promoter activity, message stability and transcriptional control. Another source of gene expression variability is plasticity, whereby an organism adjusts gene expression to alter its phenotype in response to a changing environment[6]. However, gene expression patterns can also vary among genetically identical cells in a constant environment[7–10]. This is commonly described as “noise”.

Expression variability, whatever its source, is an evolvable trait subject to natural selection, whereby each genes have an optimal expression level and variance required for an organism’s fitness and selection minimizes this variability[7, 10–14]. In this case, genes with low variability have been subjected to heavy selection pressure to minimize population expression variance. Conversely, high variability genes have been selected for high variance. Genes with high expression variability could be drivers of phenotypic diversity, as suggested by position association between expression noise and growth[15–18]. In this interpretation, genes with high variability allow for growth in fluctuating environments. Understanding the role of the gene expression variability patterns across human populations and in isogenic mice will therefore provide crucial insights into how genetic differences contribute to phenotypic diversity, susceptibility to disease[19, 20], differentiation of disease subtypes[5], development[21–24], and alterations in gene network architecture[25].

In this analysis, we used a novel method to analyze global gene expression variability in non-diseased human breast, cerebellum, and frontal cortex tissues. Our method differs from other protocols in that we account for sampling error in our analysis as well as estimate expression variability independent of expression magnitude. In addition, we analyzed gene methylation in conjunction with expression variability. Our work suggests that expression variability is an important part of the development and aging process and that identifying genes with very high or very low expression variability is one way to identify physiologically and important genes.

## Results

### Estimating expression variability

We measured human gene expression variability (EV)[1] in post-mortem non-diseased cerebellum (n = 465) and frontal cortex samples (n = 455) and biopsied normal breast tissues (n = 144). Gene expression was measured using the Illumina HumanHT-12 V3.0 expression BeadChip. We excluded probes corresponding to non-coding transcripts as well as those with missing probe coordinates, resulting in a list of 42,084 probes. We chose to estimate EV of a microarray probe independent of its expression magnitude. In this respect, neither the coefficient of variation nor variance are suitable. The former has a bias for genes with low mean expression and the latter has a bias for high mean expression genes. We modified the method initially described by Alemu et al[1]. First, we calculated the median absolute deviation (MAD) for each probe. Then we modelled the expected MAD for all probes as a function of median expression using a locally weighted polynomial regression (Fig. 1A, red line). The expected MAD regression curves for each tissue type exhibit a flat, negative parabolic shape where the lowest and highest expression probes represent the troughs of the curve. Variability in gene expression levels has previously been shown to decrease as expression approaches either extrema[7, 9, 26]. The EV for each probe was calculated as the difference between its bootstrapped MAD and the expected MAD at each median expression level (Fig. 1A). Positive EV values indicate that the probe has a greater expression variability than probes with the same expression magnitude mean. Conversely, negative EV values imply reduced population expression variability. We next plotted the kernel density estimation function of EV for each tissue (Fig. 1B). The EV distribution in all three tissue types exhibit large peaks around the zero mean and a long tail for positive EV probes. Breast tissue exhibited a larger shoulder of the negative EV probes compared to cerebellum and frontal cortex tissues. This is likely attributable to the lower number of breast samples (144 compared to 456 and 455 samples respectively).

**Figure 1.**
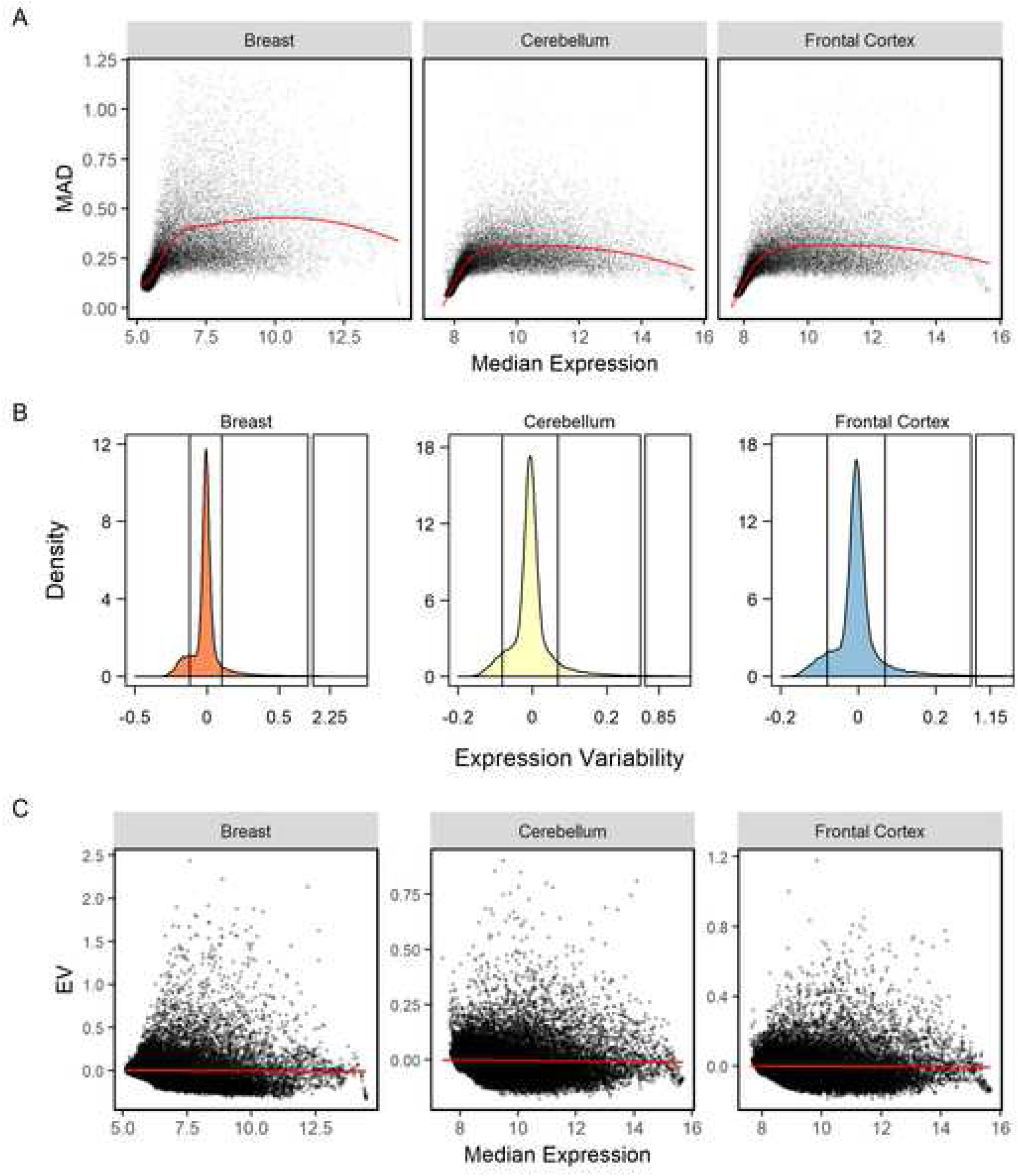
Expression variability (EV) in human breast, cerebellum, and frontal cortex tissue. (A) Expected expression MAD for curve as a function of median probe expression (solid black line). (B) Kernel density estimation function of EV. The vertical black lines represent the EV classification ranges. (C) Expression variability as a function of median gene expression. Adjusted R^2^ values for the linear regression model shown in red were 0.0002, 0.0008, and 0.005 and the associated Kendall rank correlation coefficients were ×0.208, ×0.201, ×0.213 for breast, cerebellum, and frontal cortex tissues respectively.

We then confirmed the independence of EV on expression by modelling the relationship between the two variables using a linear regression (Fig 1C) and calculating the Kendall rank correlation coefficient for each tissue type (Table 1). Based on the poor adjusted R^2^ values and Kendall rank correlation coefficients, we conclude that there is no substantial correlation between probe EV and expression magnitude.

**Table 1.** Correlation analysis of EV and probe expression. Adjusted R2 values were calculated using a linear regression model.

|  | Breast | Cerebellum | Frontal Cortex |
| --- | --- | --- | --- |
| Kendall Rank Correlation Coefficient | -0.208 | -0.201 | -0.213 |
| Linear Regression Adjusted R<sup>2</sup> Value | 2x10<sup>-4</sup> | 8x10<sup>-4</sup> | 5x10<sup>-3</sup> |

Next, we then classified each probe into three categories based on their EV. We used the term “Hyper-Variable” to describe probes whose EV was greater than 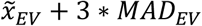. Probes with an EV less than 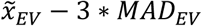 were deemed “Hypo-Variable”. The remaining probes that fell within the range of 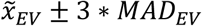 were considered “Non-Variable”. A probe classified with a “Non-Variable” EV means that its bootstrapped MAD is similar to the MAD of all genes with similar expression magnitude. It is important to note that these probes still have expression variability across the population. We propose that these three distinct groups, categorized based on EV, correspond to distinct functional and phenotypic gene characteristics.

### Statistical nature of Hyper-variability

A previously unexplored aspect of expression Hyper-variability is the statistical characteristics of expression amongst genes with this wide range of gene expression. Specifically, high EV could be the result of a multimodal distribution of gene expression with two or more distinct expression means or might simply result from a broadening of expression values around a unimodal mean value. In order to distinguish between the two possibilities, we modeled each probe expression as a mixture of two Gaussian distributions prior to estimating probe EV (Fig. 2). Next, we identified the peaks of the kernel density estimation function for each Gaussian distribution and compared the distance between the peaks as well as the ratio of peak heights. Probes with peaks that were greater than one median absolute deviation apart and displayed a peak ratio greater than 0.1 were classified as having a bimodal expression distribution. Probes that did not satisfy both criteria were considered to have a unimodal distribution. Only a small minority of the probes (16/41,968 breast tissue probes, 6/41,968 cerebellum probes, and 6/41,968 frontal cortex probes) showed a bimodal distribution of gene expression. The remaining majority of Hyper-Variable probes had a unimodal distribution. This indicates that high expression variability is a result of a widening of possible expression values across a single mean rather than the gene expression existing in two or more discrete states.

**Figure 2.**
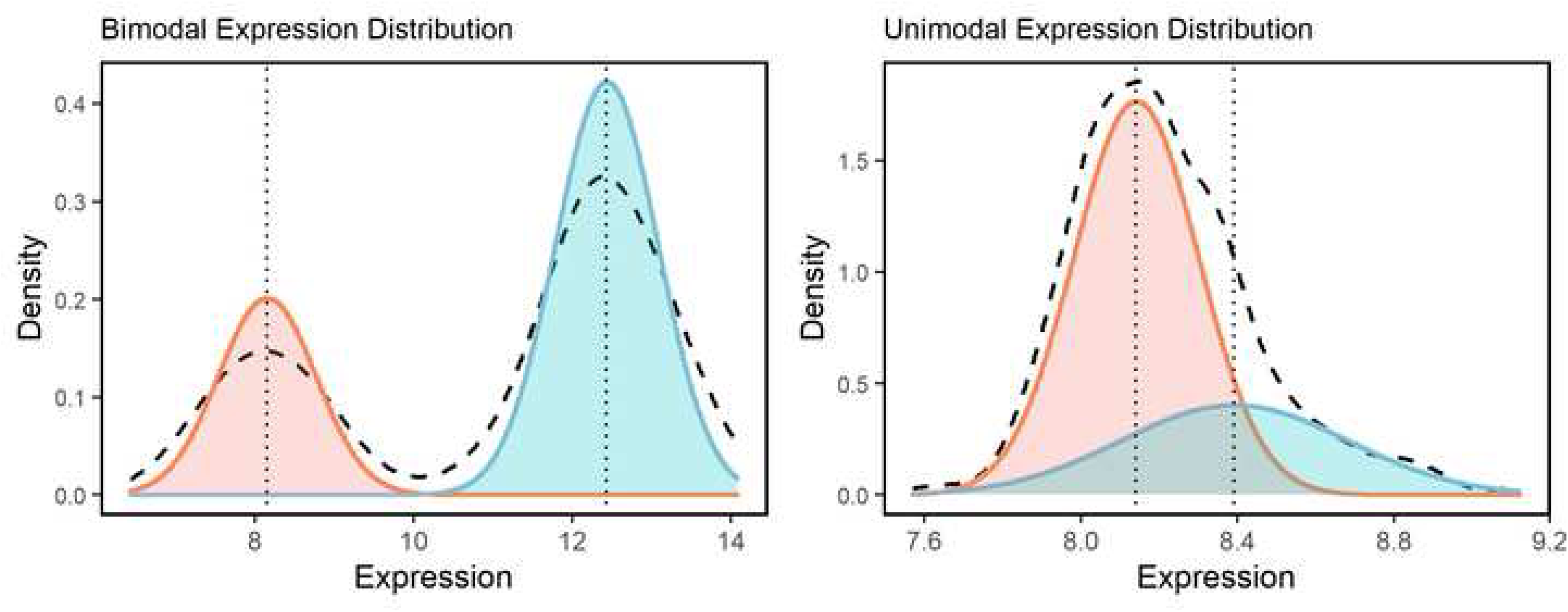
Bimodal Hyper-Variable gene expression detection. Gaussian mixture modelling method of detecting bimodal probes. The dashed lines represent the overall gene kernel density estimation function of gene expression. The two Gaussian models are shown in dark grey and light grey, and the dotted vertical lines represent the distribution means.

### Accounting for sampling error in EV classification

We were concerned that the classification of a probe into Hyper-, Hypo- and Non-Variable classes might be the result of sampling errors. To minimize this possibility and to increase the accuracy of our EV classification method, we divided each of our tissue samples into two equally sized sample probe subsets and repeated the EV analysis. This 50-50 split-retest procedure was repeated 100 times with each iterative retest using a random split of the probes. Fig 1B shows the kernel density estimation function of a concordant EV classification for each probe into Hyper-, Hypo- and Non-Variable class across the three subsets in each tissue type. Fig 3A demonstrates that classification of a probe as Hyper or Hypo-Variable based on a single analysis of the population is problematic due to sampling bias. We see a substantial decrease in the number of probes in the Hyper- and Hypo-Variable probe sets after conducting our split-retest protocol (Fig. 3B and Table 2). Thus, our split-retest method likely increases the robustness and accuracy of EV classification.

**Figure 3.**
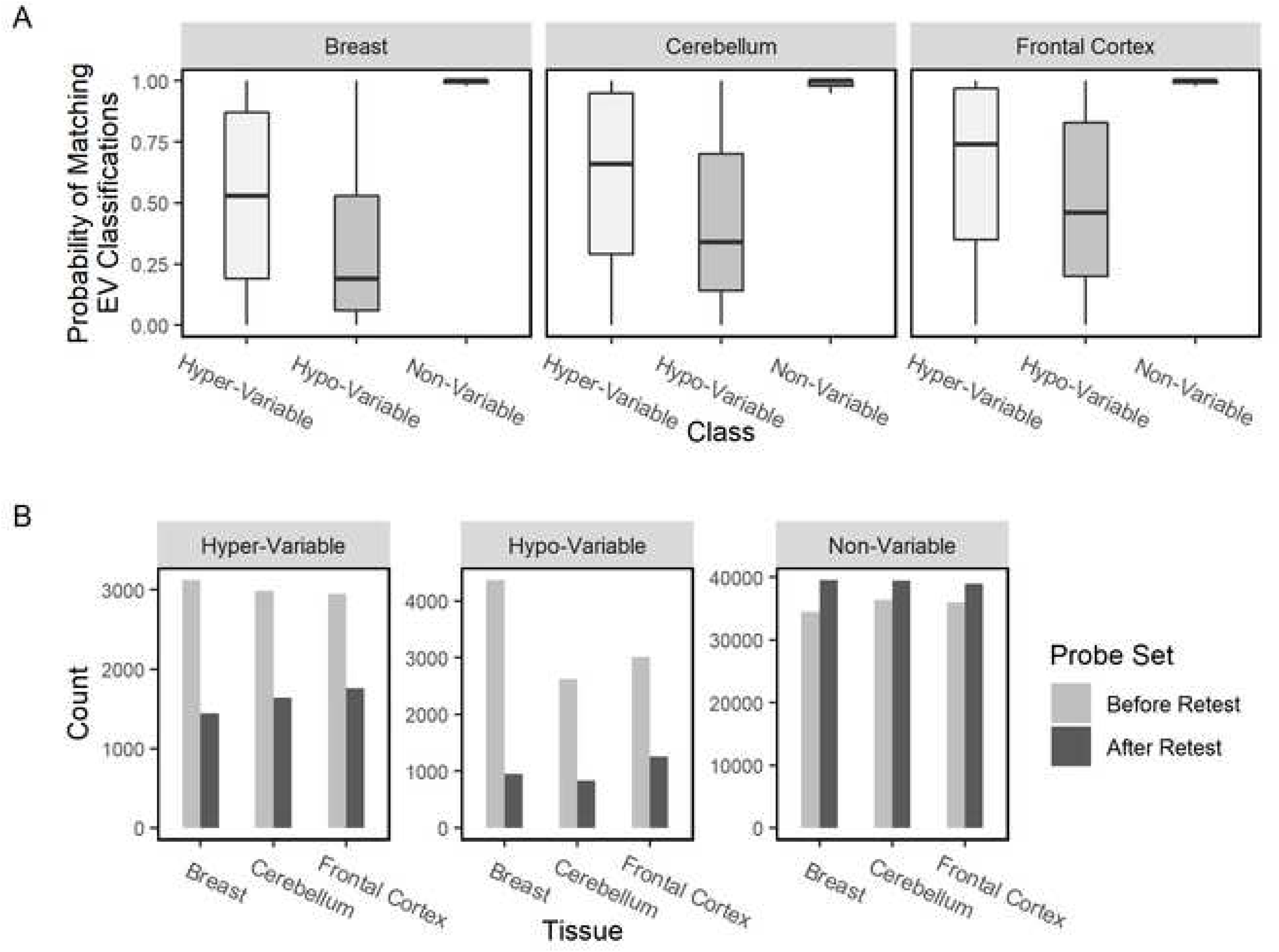
Cross-Validation of EV Classifications. (A) Relative frequency of EV classification accuracy between original distribution and 50-50 split retest replicates (n=100). (B) Number of probes in each EV probe set before and after split-retest protocol.

**Table 2.** Count summary of probes before and after 50-50 split-retest procedure. Hypervariable and Hypovariable probes that were not retained after the split-retest were relabeled as “Non-Variable”.

| Probe Set | Tissue | Number of Probes Before Retesting | Number of Probes After Retesting | % of Probes After Retesting |
| --- | --- | --- | --- | --- |
| Hypervariable | Breast | 3125 | 1448 | 46.34 |
|  | Cerebellum | 2987 | 1640 | 54.90 |
|  | Frontal | 2949 | 1760 | 59.68 |
| Hypovariable | Breast | 4371 | 957 | 21.89 |
|  | Cerebellum | 2619 | 837 | 31.96 |
|  | Frontal | 3019 | 1254 | 41.54 |
| Non-Variable | Breast | 34456 | 39547 | 114.78 |
|  | Cerebellum | 36356 | 39485 | 108.61 |
|  | Frontal | 35994 | 38948 | 108.21 |

### Tissue-specificity of EV

We next mapped Hyper-, Hypo and Non-Variable probes onto their respective genes. Individual genes can have multiple probes attached to them and we refer to the identified genes as being “probe-mapped”. A probe-mapped gene is assigned to a Variability group if one or more of its probes have that characteristic Variability. Thus, the possibility exists that an individual gene could be placed in one or more Variability groups based on differential behavior of probes mapped to that gene. However, the number of genes that have are classified in one or more Variability groups involved is small (Breast: 2.22%, Cerebellum: 2.76%, Frontal Cortex: 3.18%).

Because we have calculated EV from different tissues, we were able to determine the extent to which tissue-specific factors might contribute to EV. This is an important question because expression variability exists not only between individuals but between different tissues in the same organism.. As shown in Fig. 4A, only a small minority of Hyper-Variable and Hypo-Variable probe-mapped gene sets are shared between the three tissues. 16% of the Hyper-Variable probe-mapped genes were classified as such in the three tissues and 18-26% of the Hypo-Variable were so classified. The Non-Variable probe-mapped gene sets contained over 82% of genes in each tissue type, with over 71% of the measured genes commonly classified as NV in all three tissue types.

**Figure 4.**
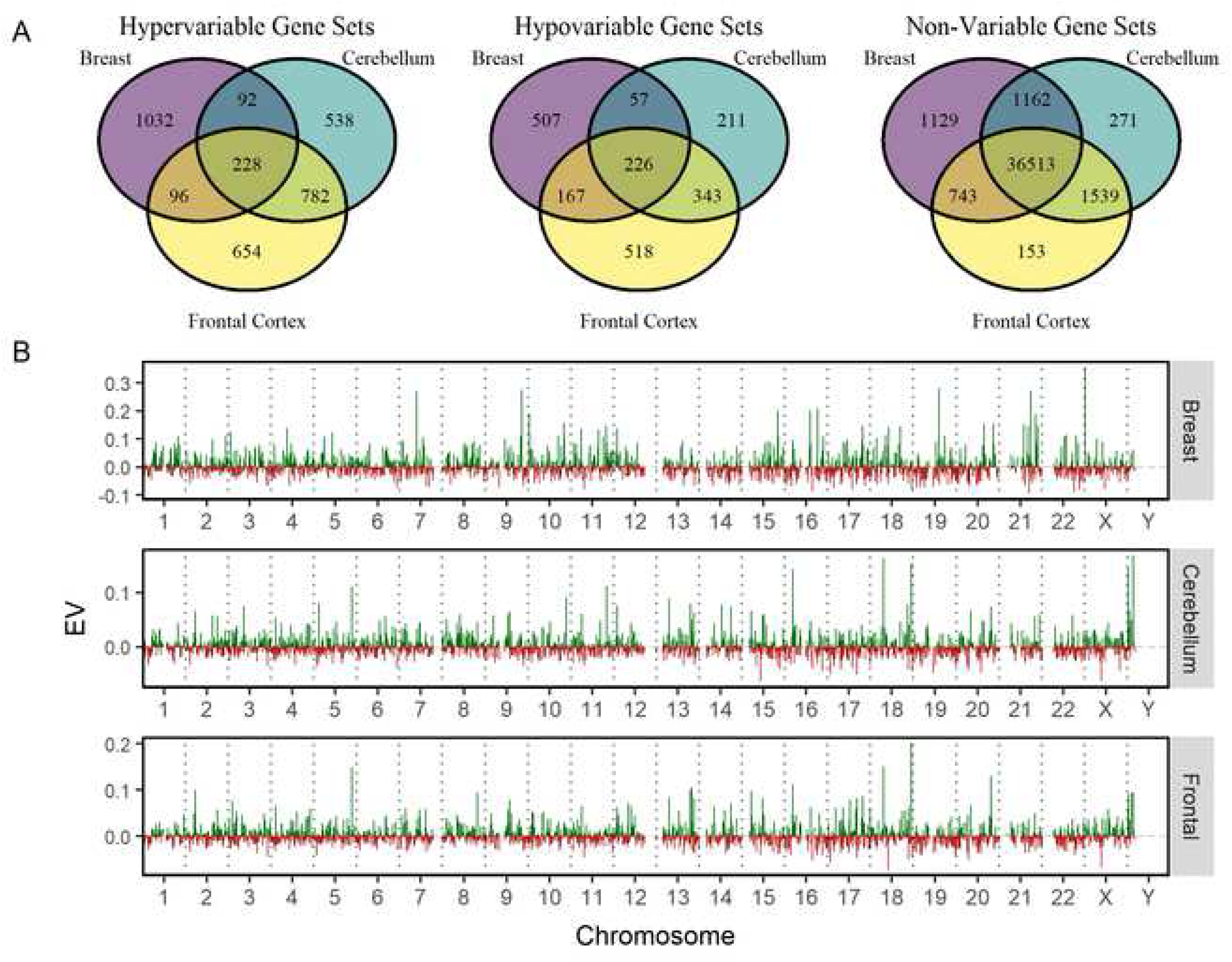
Tissue Specificity of EV. (A) Venn diagrams comparing EV classifications of probe mapped genes sets between breast, cerebellum, and frontal cortex tissues. (B) Effect of genomic position on EV. Each chromosome is divided into 100 bins (x-axis) based on the maximum gene coordinate annotation, and the average EV in each bin is measured (y-axis). Bins with an average EV greater than 0 are represented in green, while those with a negative EV are represented in red. Bins with less than three probes were assigned an average EV of zero.

### EV and gene structural characteristics

To understand possible genomic mechanisms by which population expression variability occurs, we first explored the relationship between EV and various structural features of the genes. Expression variability has previously been reported to be associated with gene size, gene structure, and surrounding regulatory elements[1]. However, we found no significant linear correlation between EV and a gene’s exon count, sequence length, transcript size, or number of isoforms (Additional file 1). While certain linear models exhibited statistical significance (p < 0.05), the fit of the model and subsequent comparison of the linear model against a local polynomial regression curve showed that the correlation was either too small to draw a conclusion or not correctly defined by a linear model.

While we did not find that the physical gene characteristics were correlated to EV, previous studies have shown that the position of a gene on a chromosome has considerable effects on stochastic gene expression variability [27]. We next tested if there is a relationship between expression variability and chromosomal position (Fig. 4B). To this end, each chromosome was divided into 100 bins and the mean EV all the genes within each bin determined. We display mean EV so that the graphed value does not depend on the probe density. However, bins that have a small number of probes may skew positional values. We therefore introduced a minimal threshold for number of probes in each bin. Any bin with less than 3 probes would be considered to have a zero EV value. We found that EV is not uniformly distributed across the genome, and individual regions of chromosomes exhibited peaks of high expression variability or troughs of low expression variability. To further confirm our conclusion, we tested the cosine similarities of the chromosomes within and across the tissue types (Additional file 2). This similarity analysis is consistent with the idea that EV is not randomly distributed throughout the genome. Furthermore, chromosomal EV distributions across chromosomes exhibited low similarities with each other. Because the probes used for the three different tissues are identical, this conclusion is not affected by probe density.

### Functional analysis of Hyper-, Hypo- and Non-Variable genes

In order to understand the overall biological significance of EV, we examined the functional aspects that are enriched in the Hyper-Variable, Hypo-Variable, and Non-Variable probe-mapped gene sets by conducting a gene set enrichment analysis in each category. We conducted a functional enrichment analyses of the gene symbols corresponding to the probes in each probe-mapped gene set. We determined the over-represented Gene Ontology (GO) terms that were unique in each tissue type, as well as GO terms that were common in all three tissue types. The resulting GO annotations were simplified and visualized using a REVIGO treemap. The top five terms for each tissue type can be found in Table 3, while the complete list of GO term treemaps can be found in Additional file 3. It should be noted that the GO term “Proteolysis involved in cellular catabolism” appears both in the “Common Probe-Mapped Genes” and “Breast-Specific Probe Mapped Genes” for the Hypo-Variable set. The genes involved in both cases are unique but they are members of the same GO pathway.

**Table 3.** Top 5 common and tissue-specific REVIGO GO annotations in the Hyper-Variable and Hypo-Variable probe mapped gene sets of breast, cerebellum, and frontal cortex tissues.

|  | **Common Probe-Mapped Genes** | **Breast-Specific Probe-Mapped Genes** | **Cerebellum-Specific Probe-Mapped Genes** | **Frontal Cortex-Specific Probe-Mapped Genes** |
| --- | --- | --- | --- | --- |
| **Hyper-Variable** | Regulation of bone remodeling | Epithelial cell differentiation | Regulation of nervous system development | Histamine secretion |
|  | Regulation of inflammatory response | Primary alcohol metabolism | Regulation of transmembrane transport | Regulation of cell morphogenesis |
|  | Response to zinc ion | Positive regulation of cellular component movement | Regulation of neuron death | Trans-synaptic signaling |
|  | Carboxylic acid biosynthesis | Response to corticosteroid | Negative regulation of response to external stimulus | Regulation of neurological system process |
|  | Regulation of ion transport | Transmembrane receptor protein tyrosine kinase signaling pathway | Response to calcium ion | Dephosphorylation |
| **Hypo-Variable** | Proteolysis involved in cellular protein catabolism | Golgi vesicle transport | DNA conformation change | ncRNA metabolism |
|  | Ribonucleoprotein complex assembly | Nucleoside monophosphate metabolism | Modification-dependent macromolecule catabolism | Response to interleukin-1 |
|  | Regulation of cellular amino acid metabolism | Proteolysis involved in cellular protein catabolism | Response to camptothecin | Regulation of enter of bacterium into host cell |
|  | Innate immune response activating cell surface receptor signaling pathway | Cellular response to nitrogen starvation | Retrograde transport, endosome to Golgi |  |
|  | Negative regulation of autophagy | Mitochondrial respiratory chain complex I assembly | Regulation of ubiquitin-protein transferase activity |  |

The breast Hyper-Variable probe-mapped gene set was uniquely enriched for epithelial cell differentiation, primary alcohol metabolism, and positive regulation of cellular component movement. The cerebellum Hyper-Variable probe-mapped gene set was uniquely enriched for regulation of nervous system development, transmembrane transport, and neuron death. The frontal cortex Hyper-Variable probe-mapped gene set was enriched for histamine secretion, regulation of cell morphogenesis, and trans-synaptic signalling. The breast, cerebellum, and frontal cortex Hyper-Variable probe-mapped gene sets were commonly enriched for regulation of tissue remodeling, inflammatory responses, and responses to inorganic substances. Of note, many of the enriched GO annotations of the Hyper-Variable genes are involved in signalling pathways.

In the case of the Hypo-Variable probe-mapped gene sets, all three tissue types were enriched for protein catabolism and metabolism, ribonucleoprotein complexes, and negative regulation of autophagy. In this respect, many of the shared Hypo-Variable genes could be considered housekeeping genes. The breast Hypo-Variable probe-mapped gene set was enriched for Golgi vesicle transport, nucleoside metabolism, and protein catabolism. The cerebellum Hypo-Variable probe-mapped gene set was enriched for DNA conformation change, modification-dependent macromolecule catabolism, and retrograde transport.

### Essentiality enrichment in variable genes

Previous studies in yeast have shown that gene expression variability is reduced in genes that are essential for survival. It is believed that evolution has selected for transcriptional networks that limit stochastic expression variation of essential genes[13]. If this were true for humans, we would expect a significant number of essential genes to exhibit Hypo-Variable expression and a depletion of essential genes within the Hyper-Variable probe sets.

In order to examine a potential correlation between expression variability and essentiality in human tissues, we first tested the independence between EV classification and annotation of human essentiality (Table 4). Essentiality annotations were obtained from the CCDS[28] and MGD[29] databases. Here, direct human orthologs of genes essential for prenatal, perinatal, or postnatal survival of mice were classified as essential. Using the Pearson’s chi-square test using the chisq.test function[30] in R for the number of essential genes in each probe set (Additional File 4), we find that that the Hypo-Variable probe-mapped gene set in breast, cerebellum, and frontal cortex tissues were significantly enriched for genes with essentiality annotation. Thus, expression variability for many essential genes is constrained in humans, likely reflecting a similar biology to essential yeast genes. However, we surprisingly observe a significant enrichment of essential genes within the Hyper-Variable probe-mapped gene sets.

**Table 4.**
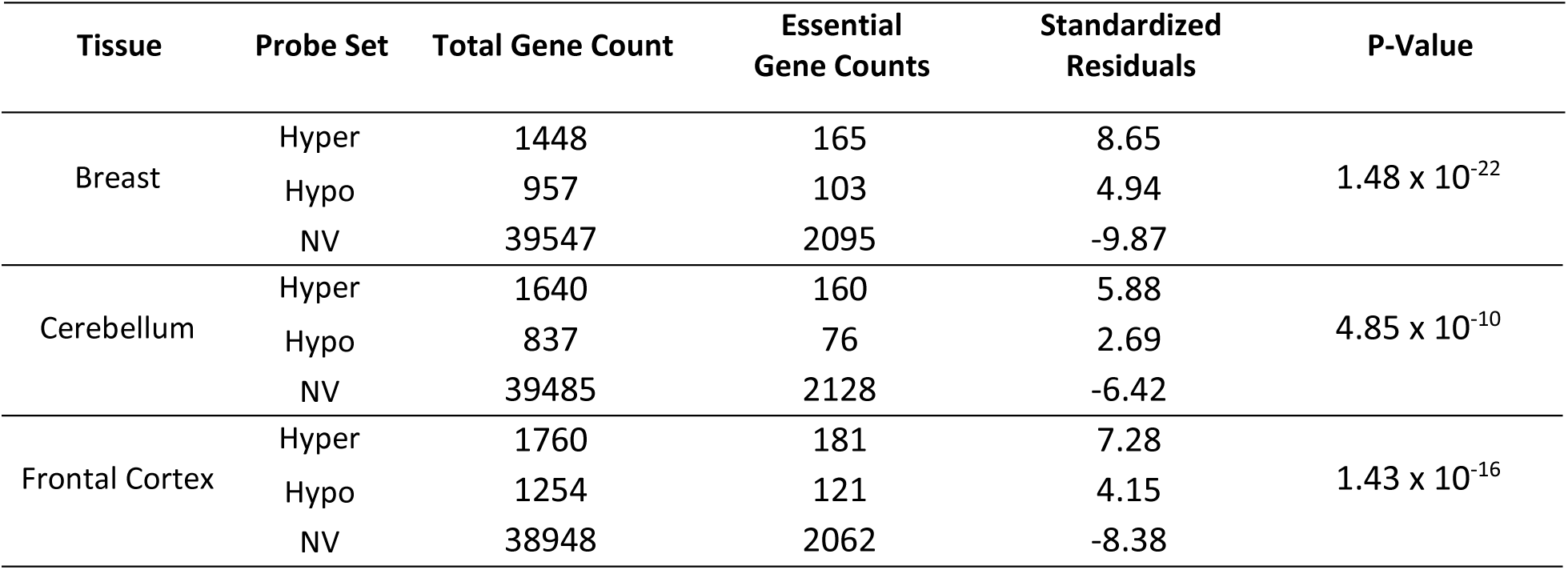
Pearson’s Chi-squared test for Essentiality in Hyper-Variable, Hypo-Variable, and Non-Variable probe mapped gene sets.

To better understand the implications of high variability in essential genes, we examined the functional annotations associated with Hyper-Variable essential genes (Table 5 and Additional file 5). The breast essential Hyper-Variable probe-mapped gene set was enriched for chordate embryonic development, cellular response to growth factor stimulus, mesenchymal cell apoptotic process, carboxylic acid biosynthesis, and cell-substrate junction assembly. The cerebellum essential Hyper-Variable probe-mapped gene set was enriched for regulation of cell development, epithelial cell migration, positive regulation of cell proliferation, cellular response to growth factor stimulus, and anterograde trans-synaptic signalling. Lastly, the frontal cortex essential Hyper-Variable probe-mapped gene set was enriched for positive regulation of cell differentiation, transmembrane receptor protein tyrosine kinase signalling pathway, epithelial cell migration, regulation of actin cytoskeleton organization, and regulation of lipase activity. Overall, the Hyper-Variable essential probe-mapped gene sets tended to be enriched for morphogenic, tissue, and organ system development.

**Table 5.** Top 5 common and unique REVIGO GO annotation subsets of Hyper-Variable and Hypo-Variable essential genes in breast, cerebellum, and frontal cortex tissues.

|  | Breast-Specific Probe-Mapped Genes | Cerebellum-Specific Probe-Mapped Genes | Frontal Cortex-Specific Probe-Mapped Genes |
| --- | --- | --- | --- |
| **Hyper-Variable Essential Genes** | Chordate embryonic development | Regulation of cell development | Positive regulation of cell differentiation |
|  | Cellular response to growth factor stimulus | Epithelial cell migration | Transmembrane receptor protein tyrosine kinase signalling pathway |
|  | Mesenchymal cell apoptotic process | Positive regulation of cell proliferation | Epithelial cell migration |
|  | Carboxylic acid biosynthesis | Cellular response to growth factor stimulus | Regulation of actin cytoskeleton organization |
|  | Cell-substrate junction assembly | Anterograde trans-synaptic signalling | Regulation of lipase activity |
| **Hypo-Variable Essential Genes** | DNA repair | DNA repair | DNA repair |
|  | Regulation of cellular protein localization | Protein oligomerization | Peptide transport |
|  | Mitochondrial genome maintenance | Positive regulation of viral process | Regulation of type I interferon production |
|  | Chordate embryonic development | Negative regulation of cell cycle | Response to UV |
|  | Protein modification by small protein removal | Lysosomal transport | Phosphorylation |

### DNA methylation and expression variability

One factor that has been postulated to regulate EV is DNA methylation. While the relationship between methylation and gene expression is complex, low promoter methylation is associated with high levels of gene expression[31–34]. Like gene expression, DNA methylation is highly variable at the cell, tissue, and individual level[35], suggesting that EV could result from variations in gene methylation. To explore this idea, we used DNA methylation annotations that were available in 724 out of 911 brain tissue samples.

DNA methylation in CpG sites is thought to be bimodal, meaning that the gene is either hypomethylated or hypermethylated[34]. In order to differentiate between low, medium, and high methylation states in our samples, we modelled gene methylation using Gaussian mixture models for the mean methylation for each gene. The distribution of gene methylation in both cerebellum and frontal cortex tissue was best modelled as a three-component system. The first component was a sub-population Gaussian mixture while the second and third components were modelled as single Gaussian distributions. Genes whose methylation fell within the first component were classified as Non-Methylated genes. Genes were classified as Medium Methylated for those in the second component and Highly Methylated if they were in third. The distribution of methylation amongst the genes is predominantly bimodal with only a minority of genes being Medium Methylated (Fig. 5A). In contrast, over 62% of cerebellum genes are non-methylated and 23% highly methylated. Similarly, 58% of frontal cortex genes are non-methylated and 22% are highly methylated).

**Figure 5.**
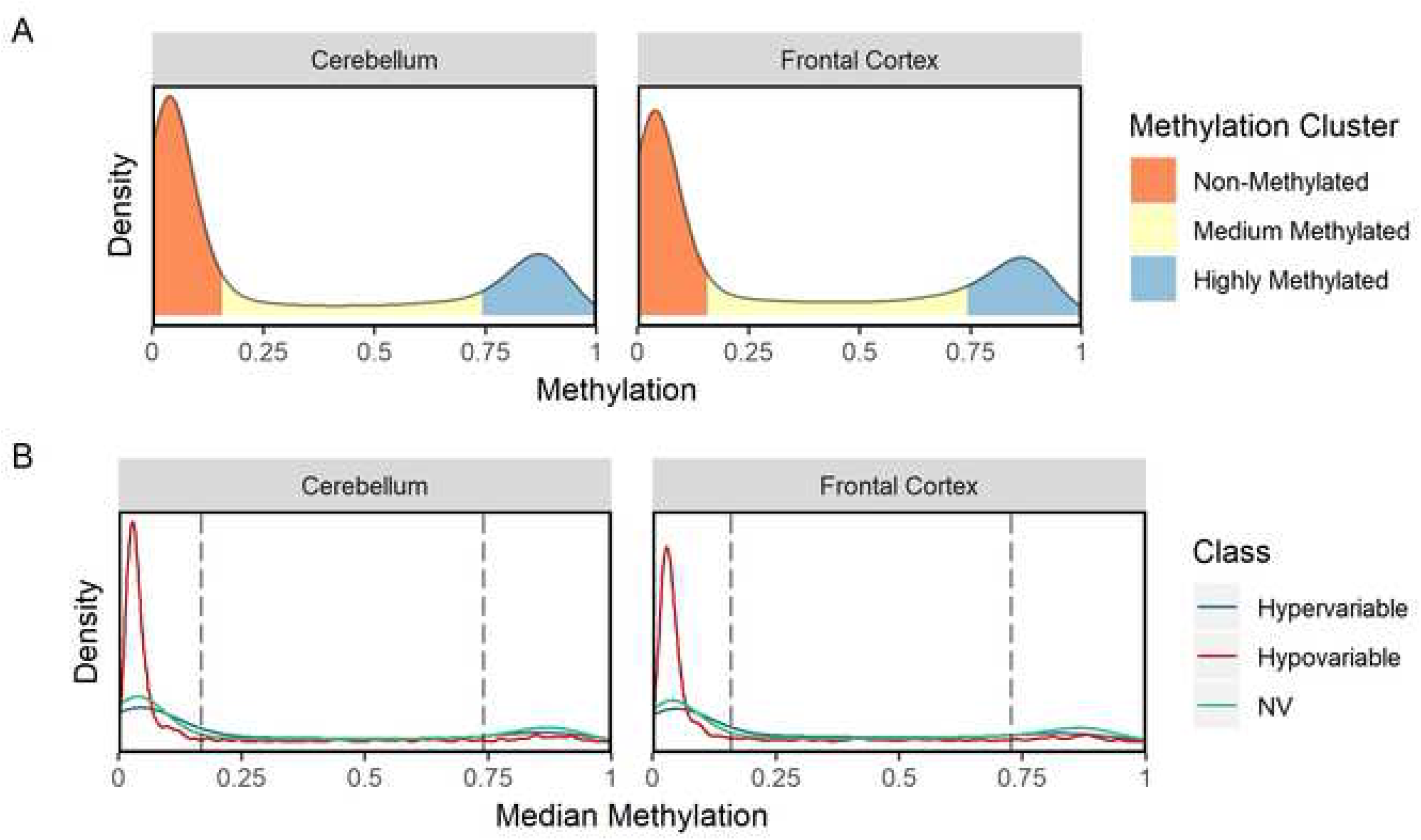
Methylation in human cerebellum and frontal cortex tissue. (A) Kernel density estimation function of average gene methylation. Gaussian mixture models were used to classify the genes into Non-, Medium- and Highly-methylated clusters. (B) Kernel density estimation function of average gene methylation by EV classification. The dashed vertical lines represent the methylation state cluster cut-offs generated by the Gaussian mixture modelling.

Next, we explored the correlation between methylation and expression based on the EV. When we subset the methylation distribution by EV classification (Fig. 5B), we observe that Hypo-Variable genes have a visibly different methylation pattern than Hyper- or Non-Variable genes insofar as Hypo-Variable genes are visibly overrepresented in the Non-Methylated gene group compared to both the Hyper-Variable and Non-Variable genes.

To further quantify the overrepresentation of Hypo-Variable genes in the Non-Methylated gene group, we conducted a chi-squared test of independence between the methylation state clusters and the EV classifications (Table 6 and Additional file 4). Both the cerebellum and frontal cortex tissues exhibited a significant relationship between the methylation clusters and EV classifications (p = 7.57 x 10^-36^ and p = 1.58 x 10^-59^, respectively). By examining the standardized residuals of the chi-square test of independence, we quantitatively confirmed the enrichment of Non-Methylated genes within the Hypo-Variable probe-mapped gene set. We also observe a significant enrichment of Highly Methylated genes in the Non-Variable gene set as well as an enrichment of Medium Methylated genes in the Hyper-Variable probe-mapped gene set. This indicates that methylation and EV classification are correlated.

**Table 6.** Pearson’s Chi-Squared Test Standardized Residuals. We tested the independence between the methylation state clusters and the EV classifications in cerebellum and frontal cortex tissues and found a significant relationship between the two variables (p = 7.57 x 10^-36^ and p = 1.58 x 10^-59^, respectively).

|  | Cerebellum Tissue |  |  | Frontal Cortex Tissue |  |  |
| --- | --- | --- | --- | --- | --- | --- |
|  | Non-Methylated | Medium Methylated | Highly Methylated | Non-Methylated | Medium Methylated | Highly Methylated |
| Hypo-Variable | 11.98 | -5.69 | -9.04 | 14.84 | -7.11 | -10.79 |
| Non-Variable | -7.52 | 0.06 | 8.59 | -10.00 | -0.04 | 11.73 |
| Hyper-Variable | 0.07 | 4.21 | -3.58 | -0.23 | 6.23 | -5.47 |

### Effects of age, sex, and PMI on variability

To further understand the biological relevance of EV, we focused on the Hyper-Variable genes to identify potential mechanisms of decreased constraint on gene expression across the samples. We systematically analyzed expression as a function of sex, age, and post-mortem interval (PMI). The breast tissue dataset lacked these clinical annotations and was excluded from this analysis. We employed a probe-wise linear regression analysis to model the relationship between Hyper-Variable probe expression and age, sex, and PMI. The resulting p-values were adjusted for multiple comparisons using the Benjamini-Hochberg procedure and considered significant when the adjusted p-value was less than 0.01. The total number of Hyper-Variable probes with sex, PMI or age as co- are shown in Table 7.

**Table 7.** Probe-Wise Multiple Linear Regression of Sex, PMI, and Age. Probes that exhibit an FDR < 0.01 are considered significant for the specific coefficient.

|  | Sex |  |  | PMI |  |  | Age |  |  |
| --- | --- | --- | --- | --- | --- | --- | --- | --- | --- |
|  | Up | Down | Total | Up | Down | Total | Up | Down | Total |
| Cerebellum | 12 | 10 | 22 | 2 | 0 | 2 | 247 | 267 | 514 |
| Frontal | 8 | 15 | 23 | 7 | 9 | 16 | 373 | 354 | 727 |

PMI might be a source of apparent expression variability because an extended PMI might compromise sample RNA integrity and lead to degradation of labile RNA[36]. Brain samples had PMI times ranging from 1 hour to 94 hours (mean = 36.14 hr), but we observe a negligible number of probes that are correlated with PMI (2 out of 1640 and 16 out of 1760 probes for cerebellum and frontal cortex, respectively). This suggests that sample integrity is unlikely to be a source of EV changes. Somewhat more surprisingly, however, is the low number of probes that are correlated with sex. Only 22 out of 1640 Hyper-Variable cerebellum probes and 23 out of 1760 Hyper-Variable frontal cortex probes show sex-dependent differences in EV. While other studies have shown widespread sex differences in post-mortem adult brain gene expression[37], EV is not substantially dependent on sex in our analysis.

However, we observe that age has a substantial effect on expression variability. Age is correlated with over 31% of Hyper-Variable cerebellum probes and over 41% of Hyper-Variable frontal cortex probes. This means that the expression of these probes becomes either more or less constrained during aging. In the cerebellum, there were 247 Hyper-Variable probes whose expression increased as a function of age and 267 genes with decreased expression. Similarly, the frontal cortex contained 373 probes with increased expression and 354 probes with reduced expression. Given that age is correlated with a considerable number of Hyper-Variable probes, we classified the age of the samples in the cerebellum and frontal cortex tissues into three age clusters according to BIC for expectation-maximization (EM) initialized by hierarchical clustering for parameterized Gaussian mixture models. The oldest cluster contained samples whose ages were between 58 and 98 (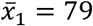). The second cluster ranged between 32 and 57 years (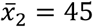), while the youngest age cluster contained samples aged 1 through 31 (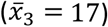).

To further explore this effect, we examined the age-dependent changes in expression of the Hyper-Variable probes across the three clusters. In each tissue type, we labeled probes whose expression was positively correlated with age as “Upregulated”, while the negatively correlated probes were termed “Downregulated”. Then, we used a hierarchical clustering method with an expression heatmap to visualize how these upregulated and downregulated probes are expressed throughout the age clusters (Fig. 6). The resulting probe hierarchical trees were clustered into groups via manual tree cutting. The complete list of GO term treemaps for significant gene clusters can be found in Additional file 6.

**Figure 6.**
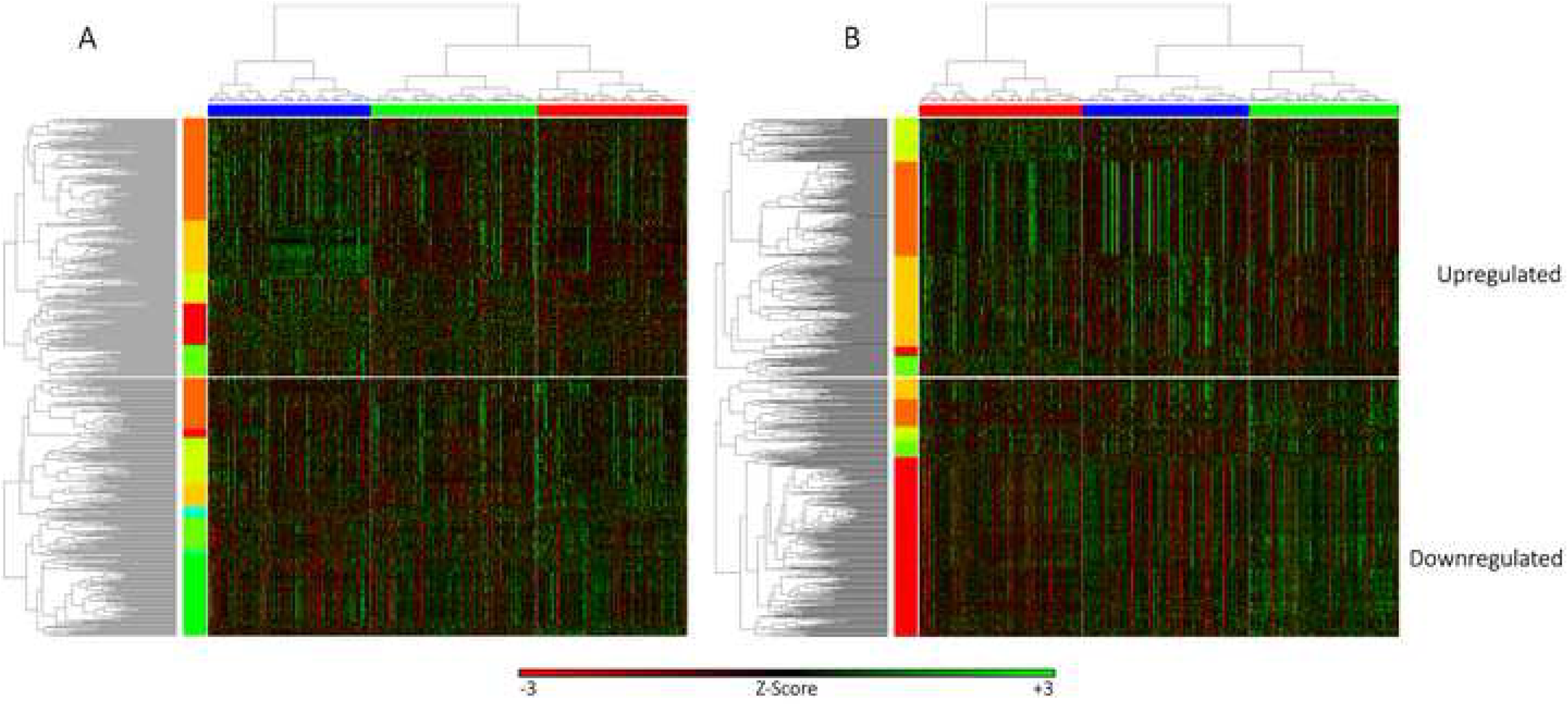
Hierarchical clustering of Hyper-Variable genes by age in (A) cerebellum tissue, and (B) frontal cortex tissue. The vertical axis represents the age-regulated Hyper-Variable genes while the samples were clustered by age and plotted on the horizontal axis. The top heatmaps represent the positively correlated age-regulated genes while the bottom heatmaps represent the negatively correlated age-regulated genes. The age clusters decrease in age from left to right in both heatmaps and correspond to the following age ranges: 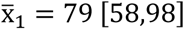, 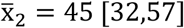, and 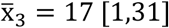.

While the cerebellum is generally considered a regulator of motor processes, it is also implicated in cognitive and non-motor functions[38]. Many of these age-dependent upregulated Hyper-Variable genes corroborate previous studies exploring the relationship between brain aging and changes in gene expression, including cellular responses to chemical stimuli (gold cluster). In particular, reactive oxygen and nitrogen species have been shown to change ion transport channel activity, and serve as an important mechanism in brain aging[39]. While all the genes selected were age-regulated, some genes exhibit outlier samples whose expression remains high across all genes in the dark orange cluster, regardless of age. These genes are more likely to be overexpressed in the samples as age increases and are enriched for peripheral nervous system neuron development and neuron apoptotic pathways. Similar enrichments of neurogenic and chemical stimuli response pathways are seen in the upregulated frontal cortex genes (gold cluster). The dark orange cluster in the upregulated frontal cortex age-dependent genes exhibits a sample-specific over- or under-expression of genes. These bimodally expressed genes are enriched for glial cell differentiation, adenosine receptor signaling pathways, and antigen processing. Lastly, we see a random scattering of expression in the yellow cluster of the frontal cortex heatmap that steadily increases with age. These genes are enriched for glial cell differentiation, cellular response to alcohol, and defense responses to fungus.

Most of the downregulated age-dependent Hyper-Variable genes in the cerebellum fall into the green cluster where expression of the genes in the cluster increases with age. These genes are involved in leukocyte-mediated immunity and defense responses to other organisms, which is supported by previous studies[40]. Interestingly, the yellow cluster exhibits U-shaped expression levels, whereby the lowest expression is seen in the middle age cluster. These genes are enriched for optic nerve development, response to interferon-gamma, and synaptic signalling. In the frontal cortex, the majority of downregulated age-dependent genes fall in the red cluster, and are enriched for ion transport, cell morphogenesis, and trans-synaptic signalling. Overall, the functional annotations of the age-regulated Hyper-Variable gene clusters suggest that population EV is one outcome of age-dependent gene expression changes.

We next investigated a possible impact of methylation status on gene expression in the Up- and Down-regulated Hyper-Variable genes. Fig. 7 shows the histogram distribution of correlation between paired gene expression and gene methylation for each gene. We observe no strong correlation between expression and methylation, suggesting age-dependent changes in expression of the age-regulated Hyper-Variable genes are not the result of methylation changes.

**Figure 7.**
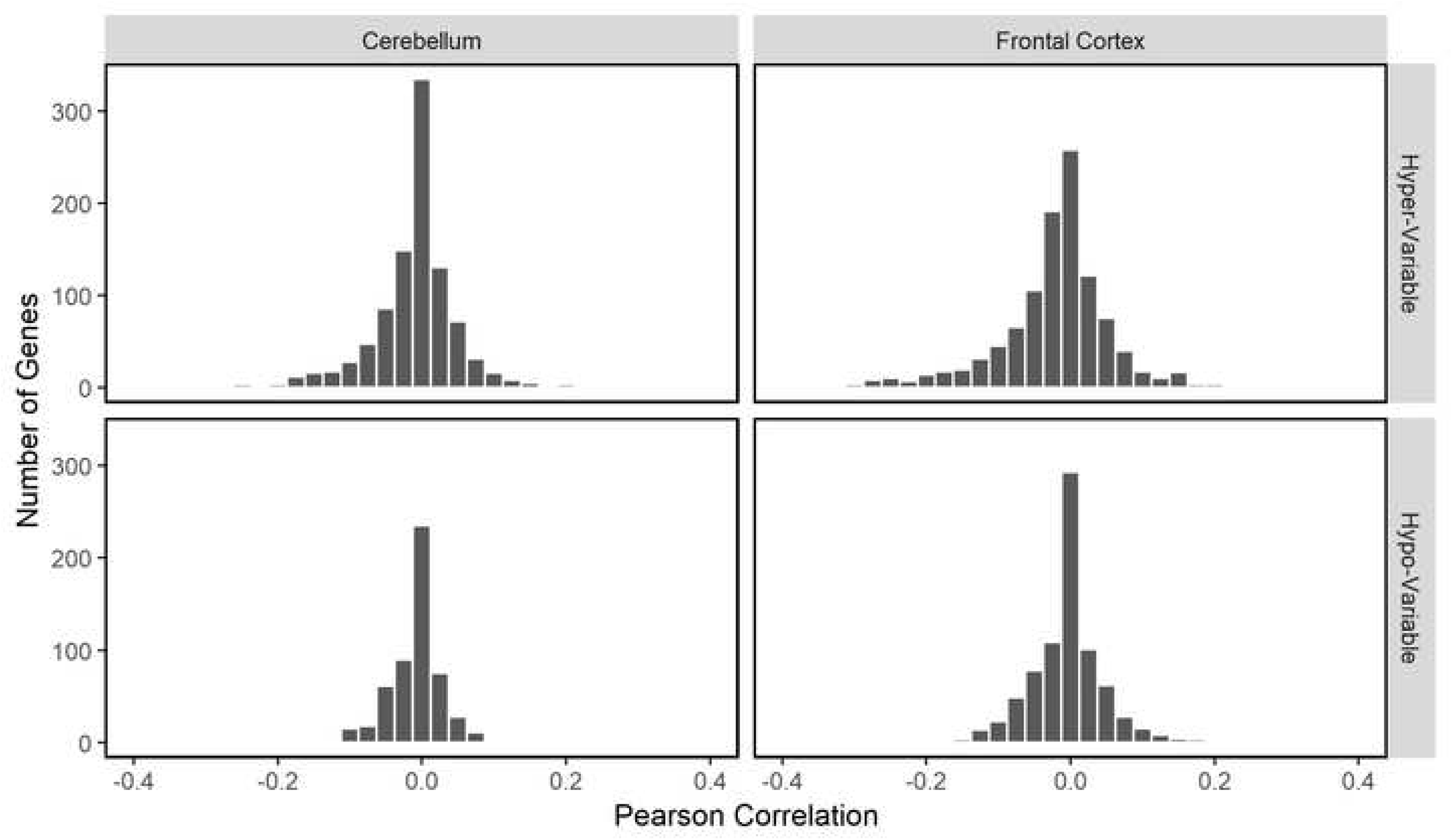
Expression and methylation correlation. Histogram of Pearson correlation coefficient between paired gene expression and gene methylation levels in the Hyper-Variable and Hypo-Variable probe sets.

## Discussion

Gene expression variability in a population is the cumulative result of intrinsic genetic factors, extrinsic environmental factors, and stochastic noise. A fundamental issue in biology is understanding the cause of expression variability within an individual organism and between isogenic and genetically dissimilar individuals of a population [42]. Expression variability has been postulated to be part of evolution, differentiation and organ homeostasis [43, 44]. In this report, we study population gene expression variability in human breast, cerebellum, and frontal cortex tissues.

Our investigation into human gene expression variability yielded several main findings. First, we find that Hyper-Variability in population gene expression is fundamentally unimodal and does not represent population switching between two or more discrete expression stages. In addition, both Hypo-Variable (highly constrained expression) and Hyper-Variable (lowly constrained expression) probe-mapped gene sets are enriched for essential genes. We observe only a small (16-26%, Figure 4A) overlap in Hyper- and Hypo-Variable probe-mapped gene sets between the three tissues, consistent with the idea that EV could be controlled by tissue-specific factors. We also find that gene methylation could have a role in expression variability. Lastly, we find that only a small number of Hyper-Variable probe-mapped genes exhibit co-variability with sex (22/1640 cerebellum probes, and 23/1760 frontal cortex probes). On the other hand, substantially more Hyper-Variable probes exhibit a strong linear association with age (514/1640 in Cerebellum and 727/1760 in Frontal Cortex).

A confounding issue with our study is the bulk nature of the tissue samples used. It is likely that multiple cell types are found in each tissue sample and that the magnitude of this heterogeneity varies between samples. This issue is not unique to our study and is common to all non-single cell sequencing studies. With respect to expression variance, cell type heterogeneity is likely to manifest itself in the identification of a gene as Hyper-Variable based on the fluctuating presence of a cell type with a unique gene expression profile. This could be one explanation for the presence of cell-type specific process in the Hyper-Variable genes associated with aging (e.g. Glial Cell Differentiation) or in the Frontal cortex-specific Hyper-Variable genes (e.g. Histamine Secretion).

However, tissue heterogeneity is only one possible explanation for Hyper-Variability. We have several reasons to suspect that tissue heterogeneity and concomitant sampling heterogeneity does not fatally compromise our analysis. First, we used large samples sizes (n>400) which would help mitigate (but not completely eliminate) heterogeneity issues. Secondly, we identified common Hyper-Variable genes between the breast, cerebellum and frontal cortex. Because of the drastic tissue type differences between these three tissues, we propose that tissue composition heterogeneity is a poor explanation for high variance gene expression common across these three tissue types. Rather, we propose that this common high variability reflects an important functional descriptor of the genes involved. Lastly, we observed that Hyper-Variable probes have an almost exclusively unimodal expression pattern (41,956/41,968 breast tissue probes, 41,962/41,968 cerebellum probes, and 41,962/41,968 frontal cortex probes). This is significant because it suggests that high EV is not the result of a chance observation of rare cell types with an unusual gene expression pattern. Nonetheless, we acknowledge that this study has not taken tissue heterogeneity into account and is a caveat to our interpretations of Hyper-Variability. Ideally, single cell analysis or sorting of the cell samples will clarify the issue. In one single cell study, Osorio et al [45] used single cell RNA-Seq to estimate gene expression variability in genetically identical human cells of three different types. Their analysis revealed that within these lines, subsets of genes with high and low expression variability could be found. They also found a positive correlation between a gene’s expression variability within a specific cell group to its variability between individuals in a population. Some genes, notably those with GO annotations for B cell activation involved in the immune response, cytokine receptor activity, cellular response to drug, and regulation of tyrosine phosphorylation of STAT protein, have a strong correlation between expression variability in single cells and in that in the population. Thus, it is likely that some of the HyperVariable genes we identified from our individuals will be genes with highly variable expression amongst cells of the same type.

On the other hand, our identification of Hypo-Variable probe-mapped genes is not affected by any potential tissue and sampling heterogeneity. These Hypo-Variable probes exhibit a restricted range of expression values in each of the samples, independent of sample heterogeneity. Shared GO annotations provided by the functional enrichment analysis of the Hypo-Variable probe-mapped genes in breast and brain tissues (Table 3) indicate that many of these genes are likely to have housekeeping functions. The definition of what constitutes a housekeeping gene is arbitrary but, in a traditional sense, it implies a strong requirement in all cell types of an organism and a limited tolerance for variations in gene expression. Some common Hypo-variable genes that would typically be considered housekeeping ones include genes for Ribonucleoprotein Complex Assembly and Regulation of Cellular Amino Acid Metabolism and Proteolysis. However, we were surprised to find a broad range of functional annotations amongst the Hypo-Variable genes. Amongst these are Negative Regulation of Autophagy, Cellular Response to Nitrogen Starvation, and Response to Interleukin-1, which would be typically be thought of as induced processes rather than obligate ones. Thus, tissues tightly regulate the expression of genes in a wide variety of processes and Hypo-Variability, similar to Hyper-Variability, is likely to be an important physiological characteristic of a gene.

The enrichment of essential genes in the Hypo-Variable probe-mapped gene sets is in agreement with previous findings in yeast showing that essential yeast genes are likely to have low expression variability. However, we detected a significant number of essential genes amongst the Hyper-Variable probe-mapped gene sets in breast, cerebellum, and frontal cortex tissue. Inactivation of these essential genes leads to pre- or neonatal fatality in mice and humans[46]. This was a surprise to us since we expected that expression of developmental genes should be tightly regulated. Our functional enrichment analysis indicates that these Hyper-Variable genes are enriched for morphogenic, tissue, and organ system development, consistent with an “essential” function yet we observe highly variable expression being tolerated. One possible explanation would be tissue heterogeneity in the samples (see above). Another possibility is these “essential” genes are required for embryonic development but have different post-embryonic roles and may not be essential postnatally. Alternatively, it is possible that these essential genes are not dose-sensitive in humans, meaning that only a certain level of baseline expression is required and expression above this baseline might be well tolerated. One additional possibility is that their protein abundance could be regulated translationally rather than transcriptionally. Inefficient translation of certain genes may have been selected for during evolution to prevent fluctuations in protein concentrations[32]. Perhaps a combination of these factors is at play.

The non-random distribution of Hyper-Variable and Hypo-Variable genes across the genome suggests that EV is dependent on epigenetic factors. Examining the methylation status of the genes allowed us to determine the relationship between gene methylation and expression variability. Firstly, we find that Non-Variable genes in the cerebellum and frontal cortex are likely to have high gene methylation. Secondly, we find that Hypo-Variable genes are likely to be non-methylated. We propose a model for methylation-dependent expression variability where the highly constrained levels of Hypo-Variable gene expression require non-methylated genes. We speculate that the lack of methylation allows transcriptional regulators requiring non-methylated DNA for binding to tightly control gene expression. On the other hand, high gene methylation reduces transcription noise and epigenetically inhibits promoter variability in human populations. Future studies should investigate the role that these putative regulators of expression play on EV, including cis-regulatory elements and transcription factors.

We find that there is limited (<26%) overlap in gene identity between Hyper- and Hypo-Variable probes in breast and brain tissue. Indeed, the chromosomal pattern of EV differs between tissue types. Our favored explanation for this is that tissue identity is created and preserved, at least in part, by changes in gene expression control pathways. Thus, genes mapped by Hypo-Variable probes in any given tissue have a constrained expression pattern because they are likely to be important in the tissue-specific function and physiology of that organelle. While there is limited overlap of genes within the corresponding EV probe-mapped gene sets of different tissues, the Hyper-Variable probe-mapped gene sets of the different tissues have similar functional enrichments and cellular protein localizations. Specifically, proteins encoded by genes mapped by Hyper-Variable probes tend to localize at the cell periphery and are enriched for cell surface signalling pathways and tissue development, including tissue remodeling and ion transport. In this respect, our work is broadly consistent with previous findings on transcript abundance in mice[23, 24]. We therefore propose that tissue identity involves high expression variability in specific tissue development pathways.

We did not observe any substantial sex dependent effects in expression variability. However, an important conclusion of our study is that many Hyper-Variable probes have age-dependent expression variability: that is, their expression significantly increases or decreases during aging. One main cause of accelerated brain aging and a causal factor of neurodegeneration is a reduction in immunological functions[47, 48]. We see evidence of downregulated immune responses in the cerebellum, specifically Leukocyte Mediated Immunity, Defense Responses to Other Organisms, and Interferon-Gamma Response pathways. Many studies also suggest that aging is associated with the upregulation of inflammatory responses[49], which is a pathogenic mechanism implicated in many age-related diseases, including cardiovascular disease, Alzheimer’s disease, and Parkinson’s disease[50]. Consistent with this idea, we see an enrichment of acute inflammatory response in the cerebellum gold cluster. Another mechanism that has been implicated with age-related diseases, such as Alzheimer’s disease and Parkinson’s disease, is synaptic dysfunction that can affect neuroendocrine signaling[51–53]. We see a downregulation of ion transport and trans-synaptic signaling in the frontal cortex, which are key components of neurotransmission and membrane excitability, and whose downregulation likely causes deficiencies in these complex processes. Furthermore, we see an upregulation of genes associated with glial cell differentiation in the frontal cortex across multiple gene clusters. Initially thought of as cells that merely support neurons, emerging research shows that neuron-astrocyte-microglia interactions are crucial for the functional organization of the brain[54]. In addition, genes specific to astrocytes and oligodendrocytes, two different types of glial cells, have been shown to shift regional expression patterns upon aging, and are better predictors of biological age than neuronal-specific genes[55]. This suggests that the Hyper-Variability and age-dependent upregulation of genes associated with glial cell differentiation or an increase in the number of glial cells in the samples.

Without examining the mechanistic control of individuals genes, it is difficult to determine if changes in gene expression result in repression or activation of their associated pathways. For example, we see an upregulation in neurogenesis-associated genes during aging in both the cerebellum and the frontal cortex, despite the common theory that neurodegeneration is a ubiquitous effect of normal brain aging. An emerging concept in neuroscience is that homeostatic plasticity of neurons is maintained through local adjustments of neural activities[56]. This overexpression of genes in pathways whose function is known to decline over time may be a compensatory mechanism for an inefficient, aging system. Within the cerebellum, a decline in neuronal function that occurs with aging may cause an upregulation of genes associated with neurogenesis pathways. In addition to mitigating neuronal dysfunction, localized increases in neurogenesis may be induced in response to cerebral diseases or acute injuries for self-repair[57]. Lastly, chronic antidepressant usage has also been shown to result in an increase in neurogenesis[58], suggesting that psychopharmaceuticals can alter neurochemistry and mimic compensatory anti-aging responses. Overall, EV plays an important role in aging, specifically in immune responses and inflammation, neurotransmission, and neurogenesis. Age-dependent gene expression could reflect a loss of regulatory control or be a part of a regulated pathway of development.

## Conclusion

Our work shows that gene expression variability in the human population is likely to be important in development, tissue-specific identity, methylation, and in aging. As such, the EV of a gene is an important feature of the gene itself. Therefore, the classification of a gene as one with Hypervariability or Hypovariability in a human population or in a specific tissue should be useful in the identification of important genes that functionally regulate development or disease. In addition, we propose that the split-retest procedure describer here is a useful technique for quantifying gene expression differences in a sample population.

## Methods

### Illumina gene expression and methylation microarray data

The analysis was conducted on two separate datasets, both utilizing the Illumina HumanHT-12 V3.0 expression BeadChip. The first dataset provides high quality RNA-derived transcriptional profiling of breast-adjacent tissue from 144 samples. The associated genotype and expression data have been deposited at the European Genome-Phenome Archive (EGA, http://www.ebi.ac.uk/ega/), which is hosted by the European Bioinformatics Institute, under accession number EGAS00000000083. The microarray readings were preprocessed using the author’s own custom script based on existing functionality within the beadarray package[59] in R and were reported as a log2 intensity. This dataset is referred to as breast tissue.

The second gene expression and the methylation datasets were catalogued by the North American Brain Expression Consortium and UK Human Brain Expression Database (UKBEC)[37, 60]. The expression data was obtained from the Gene Expression Omnibus (GEO) database[61] under accession number GSE36192. A total of 911 tissue samples were analyzed from frozen brain tissue from the cerebellum and frontal cortex from 396 subjects (Table 8). The microarray readings were processed using a cubic spline normalization method in Illumina Genome Studio Gene Expression Module v3.2.7. The expression levels were log2 transformed before any analysis. The methylation data was also obtained from GEO under accession number GSE36194. A total of 724 tissue samples were analyzed from frozen brain tissue from the cerebellum and frontal cortex from 318 subjects. The methylation microarray readings were processed using BeadStudio Methylation Module v3.2.0 with no normalization.

**Table 8.**
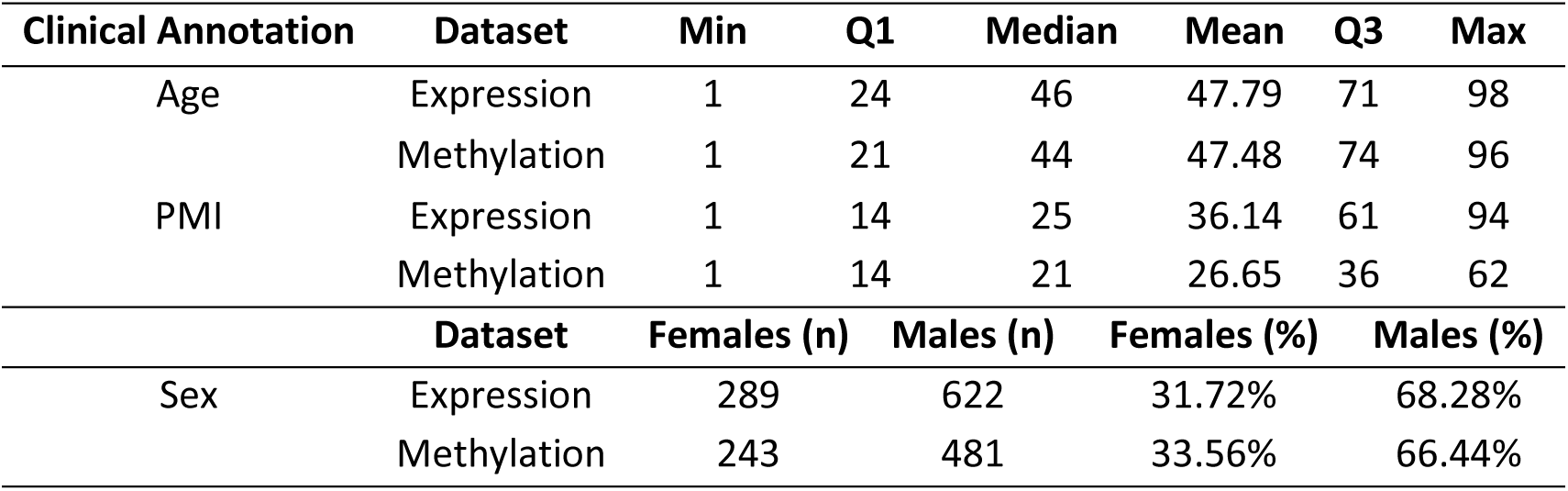
Description of brain sample dataset cohorts. Clinical annotations were not available for breast tissue samples.

### Preprocessing the datasets

Since the brain expression and methylation datasets were individually processed by different tissue banks and in several batches, we corrected for the batch effect using the limma package[62] in R. The breast tissue dataset was previously batch corrected by the authors. Next, we subset the data into groups based on the available clinical annotations provided by the NABEC/UKBEC database. These annotations included tissue type (Cerebellum and Frontal Cortex), sex (Male and Female), and age (ranging from 0 to 98 years old). We clustered the age annotations into groups using a K-Means clustering algorithm (Additional File 7), whereby the optimal number of clusters was determined using the elbow method. After four clusters, the change in total within-clusters sum of squares did not explain a significant amount of additional variance, therefore k=3 was chosen as the optimal number of clusters for the age annotation. We then converted the continuous, numeric age annotation into three categorical age groups (0-21 years, 22-73 years, 74+ years).

We then compared the 12 possible clinical annotation permutations to determine the optimal method to subset the brain samples. For each of the 12 groups, we calculated the median expression for each probe and performed a hierarchical clustering via multiscale bootstrap resampling using the pvclust package[63] in R (Additional File 7). Using an approximately unbiased (AU) p-value of 0.99, analogous to a p-value significance level of 0.01, the ideal clustering method was to subset the data solely by tissue type. Thus, we divided the brain dataset into the cerebellum tissue and frontal cortex tissue datasets. Due to the paired nature of the methylation and expression data, the methylation brain dataset was also subset into cerebellum and frontal cortex tissue subsets.

### Estimating expression variability

To calculate a magnitude-independent measure of variability for expression and methylation, we used a modified method described in Alemu et al[1]. Briefly, we first calculated a bootstrapped estimate of the median absolute deviation of each gene using 1000 bootstrap replicates. Next, a local polynomial regression curve (loess function with default parameters on R version 3.4.2) was used to determine the expected gene expression MAD as a function of the median value. No additional smoothing was used for the regression curve. We calculated gene EV as the difference between the bootstrapped MAD and the expected MAD at each gene’s median expression level.

### Identification and removal of bimodal expression probes

Probes expressions that exhibited a bimodal distribution were thought of as having two exclusive phenotypic states. However, our focus in this analysis was to examine the factors affecting the tightly regulated expression of Hypo-Variable probes or the highly variable gene expression of Hyper-Variable probes. In order to identify if a gene’s expression was unimodal or bimodal, we modeled each gene expression as a mixture of two gaussian distributions using the mixtools package[64] in R. Next, we identified the peaks of the kernel density estimation functions for each gaussian distribution and compared the distance between the peaks as well as the ratio of peak heights. Probes with peaks that were greater than one MAD apart and displayed a peak ratio greater than 0.1 were treated as having a bimodal expression and subsequently removed from the analysis. Probes that did not satisfy these criteria were considered to have a unimodal distribution and were kept for further analysis.

### EV gene set classification

We classified the probes into three distinct probes sets based on their expression variability:

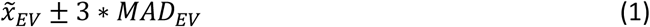

where 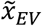 is the EV median for each dataset, and *MAD_EV_* is the bootstrapped estimate of the median absolute deviation of EV using 1000 replicates. Probes whose EV fell within the range were considered Non-Variable, those above this range termed Hyper-Variable, and the remaining were considered Hypo-Variable.

For the subsequent analyses, we used the probe sets for initial classifications then proceeded with the list of corresponding gene symbols. As such, there is a small subset of duplicate gene symbols in different EV classifications. However, the small number of duplicate genes does not significantly affect the results of the analyses.

### Bootstrapping EV gene set classifications

To statistically validate our EV classifications, we split our data into two equally sized subsets and repeated the previously explained EV method. This 50-50 split-retest procedure was repeated 100 times per tissue. Next, we determined the accuracy our of original classifications by comparing original classification of each gene with the 50-50 split classifications using a binomial test with a probability of success greater than 0.5. In this hypothesis, a “success” is defined as consistent EV classification across all three subsets, and gene classifications were considered significant with a p-value < 0.05. We also calculated the methylation variability (MV) using an identical method to EV, but did not find significant correlations between any MV classes and EV classes based on Spearman’s rank-order correlation (Additional File 8).

### Structural analysis of EV genes

Data regarding the structural features of the genes was obtained from the GRCh38/hg38 assembly of UCSC Table Browser[65]. Linear regression analyses were conducted to find any correlation between gene EV and their structural features. For the linear regression analysis of transcript size, we individually examined the largest and smallest transcripts separately. The sequence lengths excluded introns, 3’ and 5’ UTR exons, and any upstream or downstream regions.

### Gene cluster analysis

The GO term enrichment analyses were conducted using ConsensusPathDB gene set over-representation analysis[28]. The complete list of unique Illumina HumanHT-12 V3.0 expression BeadChip genes was used as a background list of genes. The resulting GO terms were then filtered manually using a q-value cutoff of 0.05. Common and unique GO terms were summarized using REVIGO[66] and visualized through treemaps by the provided R scripts. The parameters used were a medium allowed similarity (0.7) using Homo sapiens database of GO terms.

### Enrichment analyses

Using the Pearson’s chi-square test, we tested for enrichment of essential genes in each probe-mapped gene set relative to the total number of essential genes in the Illumina HumanHT-12 V3.0 expression BeadChip. A list of 20,029 protein coding genes from the CCDS database was used to test for essentiality enrichment[28]. Only genes that are solely classified as essential are considered in the analysis, resulting in a list of 2377 essential genes present in the dataset. Once the number of annotated genes and gene sets were deemed dependent variables, we determine the enrichment of annotated genes using the Pearson residuals.

The Pearson’s chi-square test was also used to test the enrichment of methylation clusters across the Hyper-Variable, Hypo-Variable, and Non-Variable probe sets.

### Classification of methylation status

In order to merge the brain expression dataset with the brain methylation dataset, we first identified the corresponding ID_REF to match the samples from each dataset. Since we could not match specific expression probe mappings to specific methylation probe mappings of CpG islands, we calculated the median probe values with a single gene mapping for both expression and methylation for each sample. This resulted in a list of median expression and median methylation of each gene for each sample. Next, we calculated the correlation between paired expression and methylation values for each gene. Lastly, we classified the genes into one of three methylation clusters based on their median methylation using Gaussian mixture models for each tissue type. In both the cerebellum and frontal cortex tissue, the distribution of median gene methylation was best modelled as a three-component system. The first component was a sub-population Gaussian mixture while the second and third components were modelled as single Gaussian distributions. Genes whose methylation fell within the first component were classified as Non-Methylated genes. Genes were classified as Medium Methylated for those in the second component and Highly Methylated if they were in third.

### Hierarchical clustering of age-dependent Hyper-Variable genes

With the exception of a few groups, the hierarchical clustering groups with the opposite sex and the same age groups tended to cluster together. While the p-values of the sex and age groupings during the hierarchical clustering were too high to warrant further subsetting of the brain dataset samples into distinct groups, they were significant enough to inspect on a gene-by-gene basis.

We used a multiple linear regression model to measure the changes in expression of the Hyper-Variable probes as a function of age, sex, and post-mortem interval (PMI):

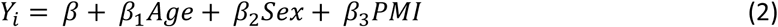

where *Y_i_* is the expression level of a probe and *β_n_* is the coefficient for each term. The p-values were calculated using a type III sum of squares regression and adjusted for multiple comparisons using the Benjamini-Hochberg method. Probes that exhibit an FDR < 0.01 were considered significant for the specific coefficient, and the sign of the coefficient determines if the probe is positively or negatively correlated with the factor.

The choice to use three age clusters as the optimal number of clusters to examine changes of EV across age samples was determined using an expectation-maximization (EM) algorithm initialized by hierarchical clustering for parameterized Gaussian mixture models in the mclust package of R. The Bayesian information criterion for each hierarchical clustering model was determined, and both the cerebellum and frontal cortex displayed identical optimal numbers of age clusters. Once the samples were correctly clustered by age, the gene clusters were selected by cutting the gene dendrograms manually. The gene expressions were then visualized as heatmaps using the gplots package[67] in R.

## List of Abbreviations

AU: Approximately Unbiased
EGA: European Genome-Phenome Archive
EM: Expectation-Maximization
EV: Expression Variability
GEO: Gene Expression Omnibus
GO: Gene Ontology
MAD: Median Absolute Deviation
MV: Methylation Variability
PMI: Post-Mortem Interval
UKBEC: UK Human Brain Expression Database

## Declarations

### Ethics approval and consent to participate

Not applicable

### Consent for publication

Not applicable

### Availability of data and material

The datasets analyzed in this study are available in the GEO repository under the accession number GSE36192 (https://www.ncbi.nlm.nih.gov/geo/query/acc.cgi?acc=GSE36192) and GSE36194 (https://www.ncbi.nlm.nih.gov/geo/query/acc.cgi?acc=GSE36194). The remaining dataset analyzed this study is available from European Genome-phenome Archive but restrictions apply to the availability of these data and are not publicly available. Data are however available from the authors upon reasonable request and with permission of EGA. All code has been deposited at https://github.com/nbashkeel/EV.

### Authors’ contributions

NB performed the computational data analysis, prepared figures, and wrote the manuscript. JL conceived, coordinated, supervised the work and helped write the manuscript. TP and MK assisted in the computational analysis and helped write the manuscript. All authors read and approved the final manuscript.

## Acknowledgements

We thank Stephane Aris-Brosou, Mathieu Lavallee, Martin Pelchat and Redaet Daniel for helpful discussion.

## Additional Files

Additional File 1: Structural analysis of genes as a function of EV

Additional File 2: EV Correlation between different tissue types

Additional File 3: Complete list of GO term treemaps for all genes

Additional File 4: Chi−Squared enrichment analysis methodology

Additional File 5: Complete list of GO term treemaps for essential genes

Additional File 6: Complete list of GO term treemaps for age-regulated Hyper-Variable genes

Additional File 7: Preprocessing of Brain Samples

Additional File 8: Methylation Variability

